# Systematic Development of Sandwich Immunoassays for the Plasma Secretome

**DOI:** 10.1101/511907

**Authors:** Ragna S. Häussler, Annika Bendes, MariaJesus Iglesias, Laura Sanchez-Rivera, Tea Dodig-Crnković, Sanna Byström, Claudia Fredolini, Elin Birgersson, Matilda Dale, Fredrik Edfors, Linn Fagerberg, Johan Rockberg, Hanna Tegel, Mathias Uhlén, Ulrika Qundos, Jochen M. Schwenk

## Abstract

The plasma proteome offers a clinically useful window into human health and disease. With recent progress made on the development of highly multiplexed immunoassays with high sample throughput, a remaining need is to establish a pipeline for validating the individual proteins that build such bio-signatures by using targeted assays. In order to streamline such efforts, we developed a workflow to build dual binder sandwich immunoassays (SIA) and chose to evaluate this on proteins predicted to be secreted form cells and tissues. Utilizing the multiplexing capacities of the bead array technology, we first screened ~ 1,800 unique antibody pairs against 209 protein targets and collected data from dilution series of recombinant proteins as well as EDTA plasma. Employing 624 unique antibodies from the Human Protein Atlas, we obtained dilution-dependent curves in plasma and concentration-dependent curves of full-length proteins for 102 (49%) of the targets. For 22 protein assays, the longitudinal, inter-individual and technical performance was determined in a set of plasma samples collected from 18 healthy subjects every third month over one year. Lastly, we compared 14 of these assays with SIAs composed of other binders, proximity extension assays and affinity-free targeted mass spectrometry. Our workflow provides a multiplexed approach to screen for SIA pairs that suggests using at least three antibodies per target. This design is applicable for a wider range of targets of the plasma proteome, while the assays can be applied for discovery but also to validate emerging candidates derived from other platforms.

## Introduction

There is a continuously great interest in increasing our understanding about those proteins that are expected to be present in blood and found outside the intracellular space, and to apply appropriate tools to discover and validate these in a given study context[1]. Such efforts preferably target the proteins that are actively secreted in comparison to those that appear in blood due to leakage, cell death or cellular turnaround. Today, the human secretome can be defined by bioinformatics tools annotating our genome based on sequences found in the protein-encoding regions[2]. Using an updated annotation[3], more than 2,600 proteins were defined as the secretome. Of these only about 600 proteins are predicted to be actively secreted to the blood while another 1,000 proteins were localized to the membrane and/or the intracellular space[3].

The technically measurable content of the human plasma proteome currently contains nearly 5,000 proteins when combining the efforts conducted with mass spectrometry (MS) as well as immunoassay platforms[4]. It has been shown that while untargeted and MS-based approaches contribute to this list with primarily cellular components, immunoassays are often more sensitive to detect low abundant proteins linked to cytokines and inflammation processes. However, and upon excluding the recent large-scale aptamer studies, only about a third of the currently annotated 2,600 proteins of the secretome[3] can be measured in plasma using other methods. Plasma profiling efforts using shotgun MS, such as those by Mann and co-workers, detected 1,200 proteins[5] in plasma. The latest versions of multiplexed immunoassay, not included in the above stated collection of plasma proteins, used 5,000 aptamers to profile 5,000 donors, as shown by Emilsson et al[6].

Here we present a complementary approach to multiplexed assays systems and systematically build sandwich immunoassays for the proteins of the plasma secretome. Our efforts are centered around the feasibility of screening and validating the antibody pairs for a variety of proteins at the same time, rather than focusing on only a few shortlisted targets. Hence it expands on previous workflows that primarily work on selected candidates[7]. Our approach was accelerated by accessing a large resource of antibodies from the Human Protein Atlas (HPA) and full-length proteins generated within the Human Secretome Project[3] (HSP) within the Wallenberg Centre for Protein Research (WCPR). The study was conducted on a multiplexed bead array platform and combined the assessment of antibody pairs using both recombinant proteins and EDTA plasma. We did not preselect the secreted proteins based on other prior knowledge or particular interest but rather availability of reagents to conduct this proof-of-concept study from screening, via validation to the analysis of longitudinal samples.

## Results

We aimed to develop a multiplexed workflow (Figure 1) to search and select for antibody pairs for the analysis of proteins secreted into human plasma (Figure 2). We combined the capabilities of the suspension bead array (SBA) technology with the resource of HSP’s full-length proteins and antibodies generated by the HPA project and investigated > 200 proteins as well as ~ 1,800 possible antibody pairs. The project was designed to be conducted in the following stages: (1) screening for possible antibody (Ab) pairs in dilution series of protein and plasma samples, (2) pre-selection of suitable pairs after assessing their apparent functionality, (3) annotation of pre-selected Ab pairs according to their binding area, (4) selection of antibody pairs for further investigations focusing on technical aspects, (5) preparation of duplex sets for plasma analysis, (6) quantification of plasma protein levels in a longitudinal sample set, and (7) compare these results with data from independent, orthogonal methods.

**Figure 1.**
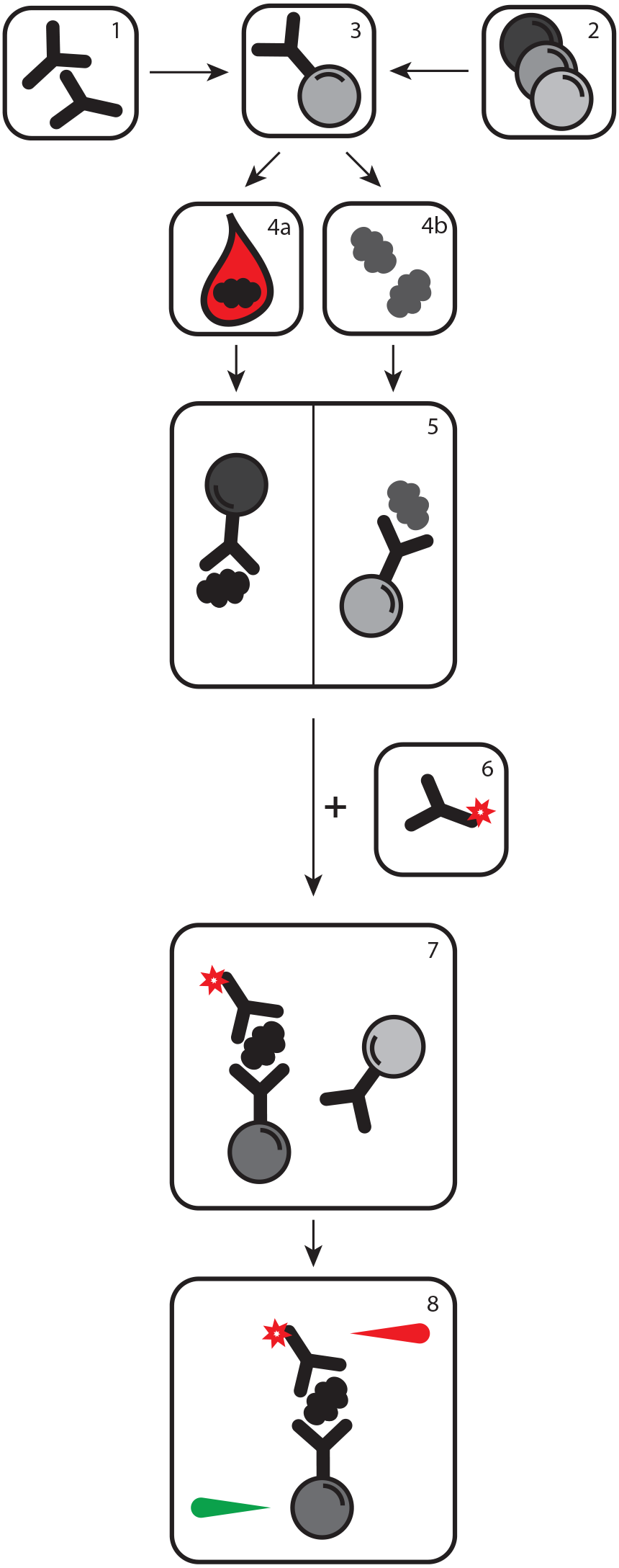
Setup for screening for SIA pairs. For capturing, Abs (1) were immobilized onto magnetic color-coded beads (2) and combined into sets of suspension bead arrays (3). A dilution series of EDTA plasma (4a) and protein standard (4b) was performed. Beads were then combined with either EDTA plasma or protein standard (5). After washing off unbound proteins, the captured proteins were detected via biotinylated antibodies (6-7). The readout occurred by the addition of a streptavidin-fluorophore and using the Luminex systems (8).

**Figure 2.**
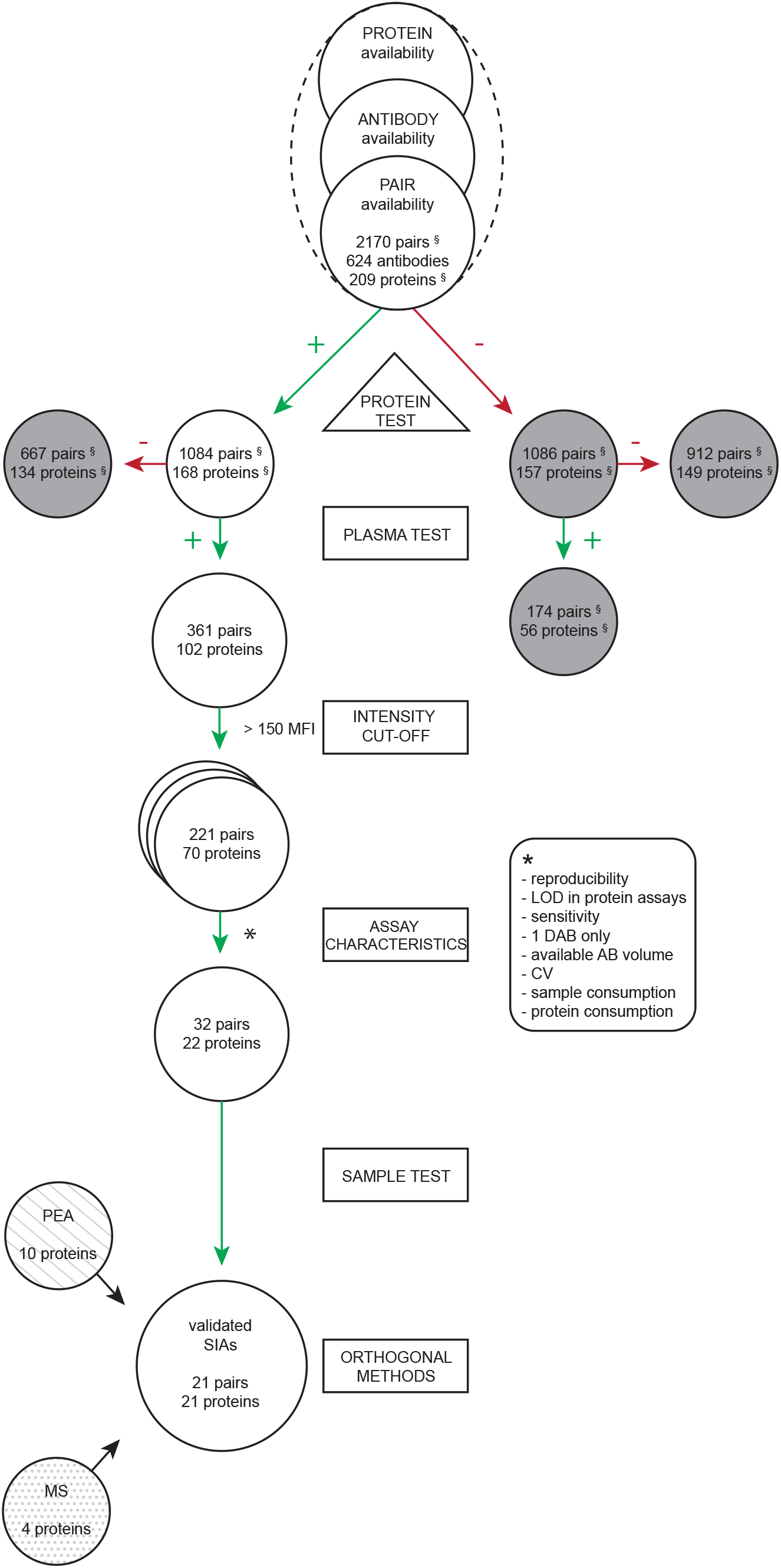
Workflow. A set of 209 protein targets, covered by 624 antibodies and resulting in 2,170 corresponding antibody pairs were selected and screened on dilution series of both, recombinant proteins as well as an EDTA plasma pool. For some proteins, more than one protein construct was tested, while some antibody pairs were duplicated (§ numbers include those). All pairs were assessed manually for concentration dependent curves in both plasma and protein. 1,084 pairs (= 168 proteins) showed a concentration dependent curve for the protein standard of which 361 unique pairs additionally detected protein in a concentration dependent manner in plasma. For these 361 pairs, corresponding to 102 unique proteins, an additional signal intensity cut-off criteria of > 150 MFI was implemented. Out of the initial 361 pairs, we used 221 for further studies in triplicates, which corresponded to 70 out of the 102 initial proteins. Results of the triplicate measurements were assessed, implementing technical aspects for exclusion, (e.g. assay reproducibility, LOD in protein assays, sensitivity towards the target protein in plasma, CV), but also additional criteria with respect to the available antibody volume, sample and protein consumption. By this 32 pairs targeting 22 proteins were left for further analysis. One pair per protein was chosen for validation according to LLOQ, using the ED50 point as the optimal sample dilution point. Lastly the selected 22 pairs were applied as SIAs for the determination of protein levels on a longitudinal plasma sample set (n = 72). For 21 pairs a protein quantification was possible, of which 14 could be compared orthogonally with data from targeted plasma mass spectrometry analysis (MS) or solution-based proximity extension assays (PEA).

### Screening for antibody pairs

#### Experimental study design

We studied a total of 209 full length proteins and used a pool of EDTA plasma samples to determine and to develop SIAs. The screening was conducted in two rounds of 109 and 124 proteins, where we aimed at replicating the findings from the first round by also including all targets with an apparent functional antibody pair, corresponding to 23 proteins, in screening round 2. For finding antibodies from the HPA resource, we chose a concentration cut-off of 0. 05 mg/ml for protein capture, and found 624 antibodies for all proteins (see Supplementary Table 1). This meant that assays could be developed using an average of 3-4 antibodies per protein, and the coverage ranged from 1-8 HPA antibodies per protein (see Supplementary Figure 2). We chose to combine 49-88 different Abs in one SBA. Among all 624 Abs, we selected those with concentrations ≥ 0.1 mg/ml and an available volume ≥ 0.5 ml as detection agents. This means that an average of 2-3 Abs were biotinylated per target protein, and the coverage ranged from 1-7 detection Abs (detAbs). Hence a total of 2,170 antibody pairs were investigated, of which 1,791 were unique.

The screening rounds were conducted by grouping the proteins into sets of 6 per assay batch and SBA. Each protein assay contained an SBA of the corresponding Abs as well as those targeting the other five proteins. Each SBA also contained control beads to judge the unspecific binding to the beads, and was distributed into 2x 384 well plates. The total number of assays per protein was defined by the number of available detAbs, and each detAb was used in 8 concentration levels of proteins and 8 dilutions steps for EDTA plasma. In total, we conducted 9,264 assays and generated 553,056 data points for the 209 proteins.

#### Reproducibility of screening results

To assess the reproducibility between the two performed screening rounds, 23 proteins corresponding to 200 unique and target matched antibody pairs were included in the second screenings. The assay conditions were in terms of dilution factor, number of dilution steps and the starting concentration for both plasma and protein maintained in the second screening. Out of the 23 targets, assay pairs for 16 targets (= 70%) revealed reproducible binding curves for both protein and EDTA plasma, as exemplified by ANGPTL3 shown in Figure 3. For each target, at least one unique antibody pair had a correlation of R^2^ > 0.92 for the protein as well as R^2^ > 0.86 in plasma. In total, 73% of all overlapping pairs had R^2^ > 0.95 in protein assays, while 59% were > 0.95 in assays with plasma. The pairs towards the additional 6 targets were regarded as reproduced for the protein assay, however the detectability in plasma was lower compared to the first screening. A higher degree of reproducibility was observed for targets that provided signal intensity levels further away from the apparent limit of detection (LOD). Additionally, using two different batches of EDTA plasma pool for the two screening rounds could have influenced the reproducibility.

**Figure 3.**
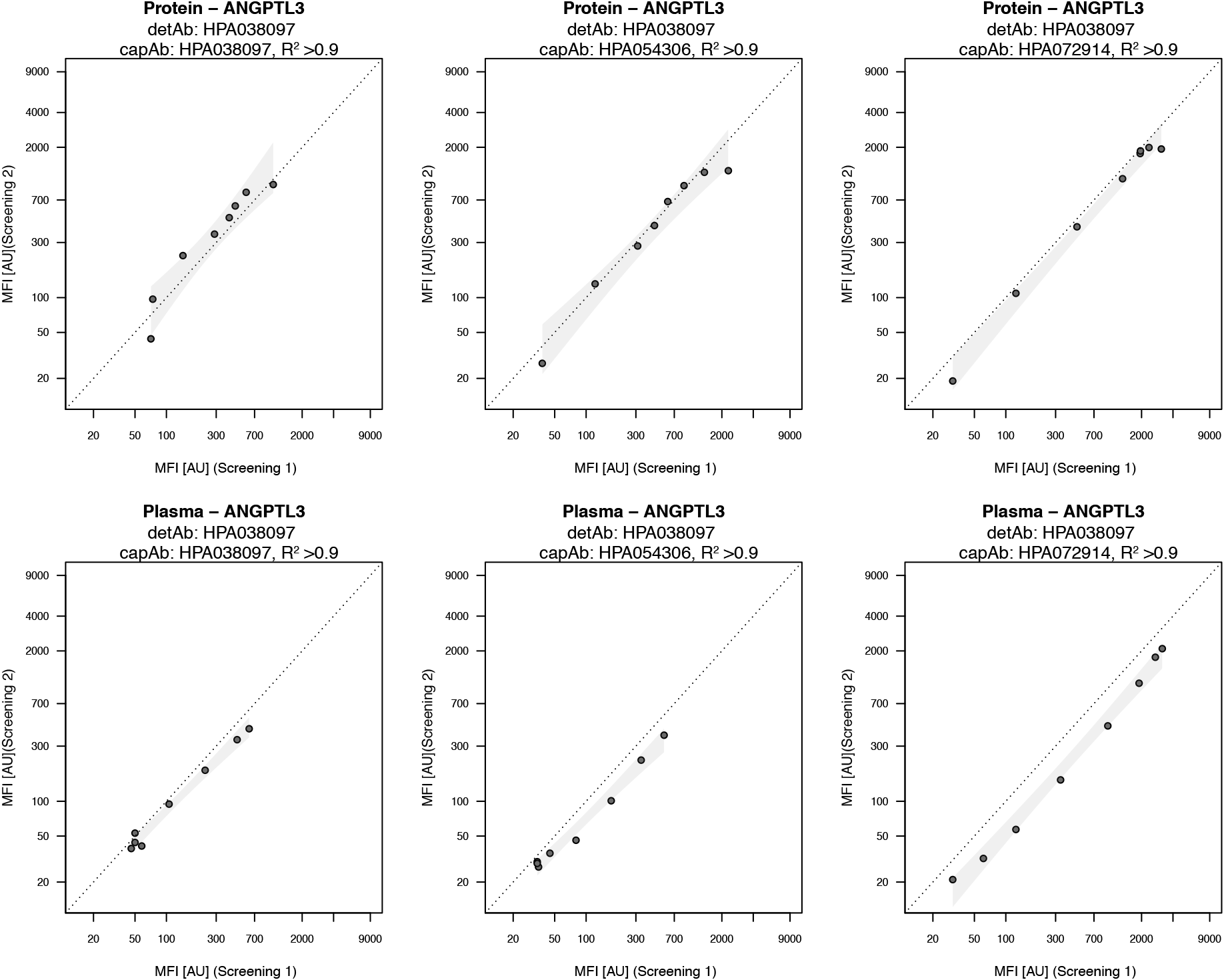
Correlation for ANGPTL3 (detAb: HPA038097) in screening assays. For each detAb correlation for all capAbs was performed both for protein standard (top) and plasma (bottom) dilution. Correlation between overlapping targets of the two screening rounds was calculated using log10 transformed MFI data and Pearson correlation with R^2^ values. Confidence interval for each pair was calculated in R based on a linear model and highlighted in the plots.

#### Pre-selection of antibody pairs

From the generated data, binding curves were manually annotated for plasma and protein in order to classify each antibody pair and select those for further optimization as illustrated in Figure 2. A summary of the outcome of the selection process is shown in Table 1. To select pairs based on their apparent functionality, we assessed the shape and concentration dependency of the curve for the expected antibody pair with the used assay conditions. We also considered all other Abs (off-target antibody pairs) included in each SBA as a background measure and noted if unexpected pairs were found in either the protein or plasma samples. All possible pairs were then assigned to one of the following four classifications according to their functionality:

1. Dilution and concentration dependent curves plasma and protein, respectively
2. Concentration dependent curves with protein only
3. Dilution dependent curves with plasma only
4. No dilution or concentration dependent curves

**Table 1.**
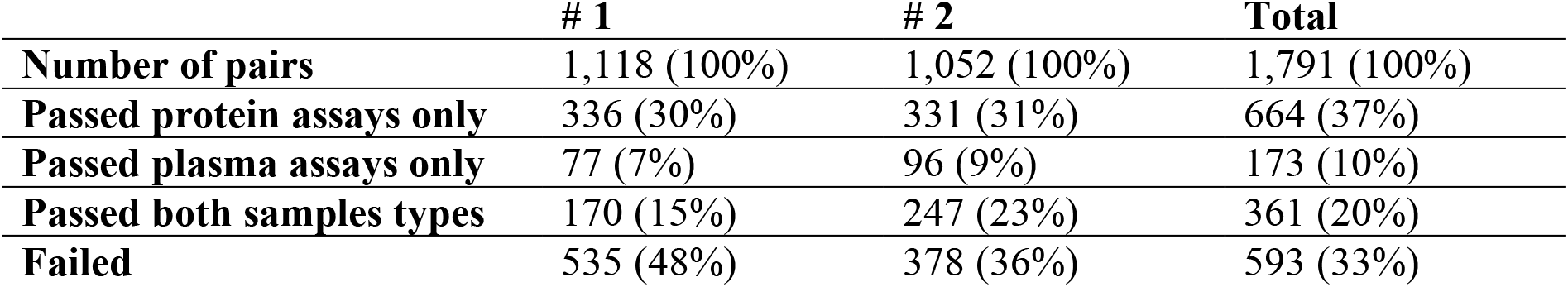
Annotation of antibody pairs determined during the screening rounds.

|  | **# 1** | **# 2** | **Total** |
| --- | --- | --- | --- |
| **Number of pairs** | 1,118 (100%) | 1,052 (100%) | 1,791 (100%) |
| **Passed protein assays only** | 336 (30%) | 331 (31%) | 664 (37%) |
| **Passed plasma assays only** | 77 (7%) | 96 (9%) | 173 (10%) |
| **Passed both samples types** | 170 (15%) | 247 (23%) | 361 (20%) |
| **Failed** | 535 (48%) | 378 (36%) | 593 (33%) |

As also summarized in Supplementary Figure 3, from almost 1,800 possible Ab pairs there were 170 from screening #1 and 247 from screening #2 that detected their target in plasma and recombinant protein. In total, 361 unique pairs were consequently annotated as “passed” and considered for further assessment and optimization. The remaining pairs did not show a concentration dependent curve for both sample types and may require further time to develop, hence were not considered for further sample analysis.

#### Annotation of pre-selected antibody pairs

As an additional assessment, we investigated the location on the protein to which the pairs of capture antibody (capAb) and detAb bound their respective target. We chose to approximate the binding areas of the Abs by using their immunogens aminoacidic sequences (22-151 residues in length) and mapping these to the sequences of the canonical protein. Here, we segmented each protein sequence into three equally long parts (N-terminal, middle and C-terminal). As shown in Table 2, there were generally more pairs for constellations that targeted the same region, which was due to using antibodies for capture and detection, generated towards an epitope located in the same region. Also, there were more pairs targeting the middle and C-terminal region than in combination with N-terminal binders. Considering the success rate for building SIAs from the screening assessment criteria, we found that on average about 12% of all pre-selected pairs passed these. A slightly higher success rate of 16% was found for purely N-terminal targeted antibody pairs as well as pairs built with a capAb and detAb targeting the C-terminal and middle, respectively (18%). The lowest success rate of ~ 6% was related to pre-selected binder pairs targeting each one of the termini. Out of the total 1791 pairs, there were 138 pairs passing the selection process of which both Abs targeted the same region.

**Table 2.** Distribution of binding regions.

| **Capture region** | **Detection region** | **Tested Ab pairs** | **% of tested** | **Passed Ab pairs** | **% of passed** |
| --- | --- | --- | --- | --- | --- |
| N-terminal | N-terminal | 176 | 10% | 29 | 16% |
| N-terminal | middle | 77 | 4% | 10 | 13% |
| N-terminal | C-terminal | 104 | 6% | 6 | 6% |
| middle | N-terminal | 85 | 5% | 12 | 14% |
| middle | middle | 434 | 24% | 48 | 11% |
| middle | C-terminal | 179 | 10% | 21 | 12% |
| C-terminal | N-terminal | 98 | 5% | 7 | 7% |
| C-terminal | middle | 165 | 9% | 29 | 18% |
| C-terminal | C-terminal | 473 | 26% | 61 | 13% |

#### Selection of antibody pairs

In a third step, we aimed at further shortlisting those pre-selected Ab pairs. As a primary criterion, the generated level of intensity (reported as MFI) was chosen as an additional cut-off in order to report signals that were > 5x above an average background level determined by the assay controls (MFI = 30 AU). Those pairs that did not reach a maximal MFI from the protein assay curves of MFI > 150 were therefore excluded. Out of the initial 361 pairs, we used 221 for further studies, which corresponded to 70 out of the 102 initial proteins.

The subsequent investigations focused on technical aspects such assay reproducibility, the apparent LOD using proteins assays, as well as sensitivity of detecting the target protein in plasma samples. All analyses were conducted using triplicates of protein concentration series.

To further resemble a sample matrix of higher complexity, the buffer used for technical assessment of protein assays and plasma samples was supplemented by adding 1% BSA. For each protein, one detAb was prioritized to limit the number of total assays. In cases where several detAbs were available after pre-selection, additional criteria for prioritization were applied: Ab pairs with the widest range of detectable concentrations of proteins in buffer and plasma, an overall lower background level in antigen-free samples, and no previous indications about possible interferences or off-target recognition of other captured proteins. The latter was possible to be observed during the screening phase, where five other proteins were also tested in parallel, as each SBA was built with a common set of 49-88 Abs covering six proteins. For the detAbs, any concentration dependent binding for other Ab-coupled beads in the SBA, such as the internal controls, were added as exclusion criteria. Lastly, the available antibody volume was considered for the polyclonal binders.

For the selection processes of Ab pairs, target proteins were regrouped into new sets of five protein targets. Concentration of the proteins and the dilution of EDTA plasma were adapted for each individual target according to the data obtained during the screening. Each assay therefore covered a broader range of concentrations in order to determine the optimal dilution point for plasma analysis. Exemplified for SIA pairs targeting ANGPTL3, CHIT1, CPA1 and FGF21, shown in Figure 4A-D, protein detection was specific and accompanied by only very minor increase in signals from other beads. When analyzing plasma, we found that background signals from other antibodies arose when using plasma at a lower dilution than 1:50 dilutions. Still, at a plasma dilution of 1:12, the intended signals were > 5-fold above any other binder pair.

**Figure 4A-D.**
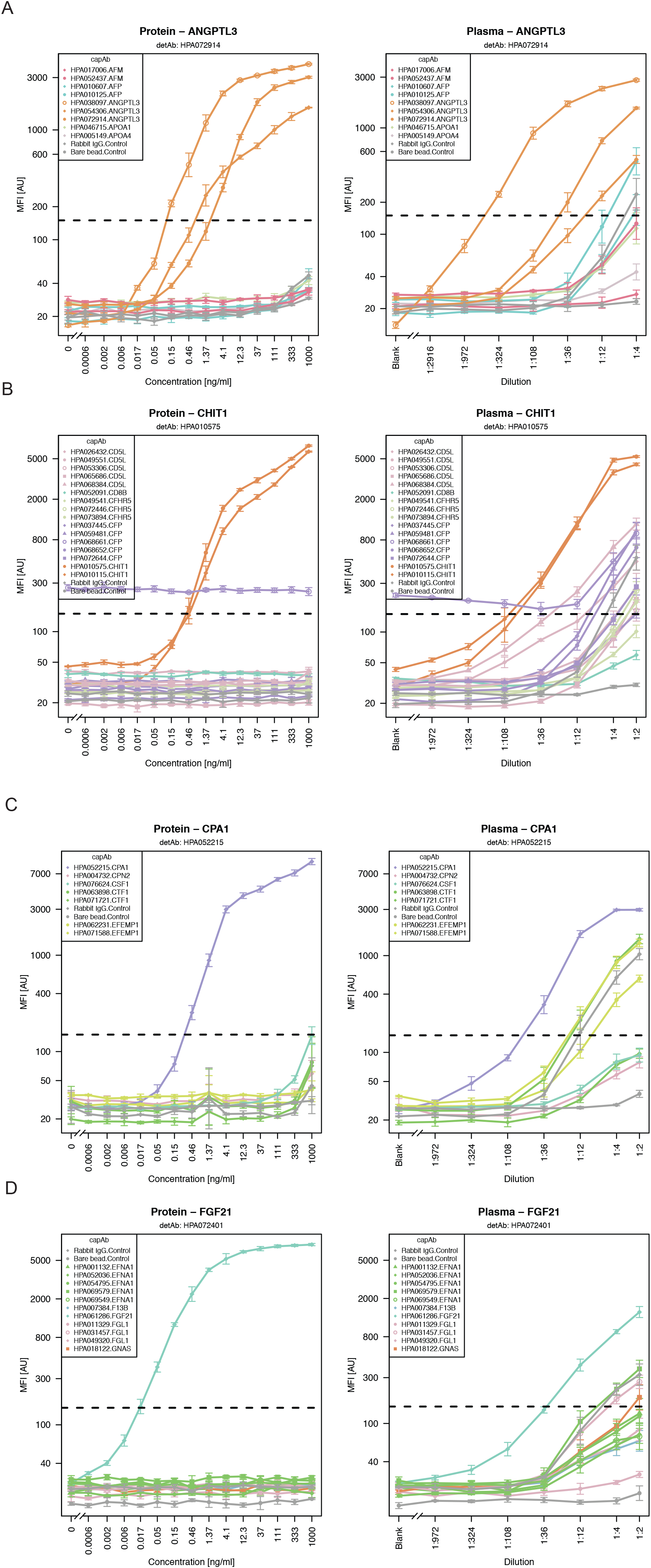
Dilution curves during selection phase for ANGPTL3, CHIT1, CPA1 and FGF21 in protein standard and plasma. During selection phase, dilution curves for all targets including all capAbs corresponding to one detAb were plotted to evaluate the performance of the different pairs. The signals above background (> 150 MFI) are indicated by a horizontal dotted line. Exemplary dilution curves for ANGPTL3 (A), CHIT1 (B), CPA1 (C) and FGF21 (D) in recombinant protein standard (left) and EDTA plasma (right) are shown.

Out of the 221 pairs targeting 70 proteins, we found 43 pairs for 27 proteins suitable for further analysis according to the criteria stated above. To further find the best performing Ab pairs for one protein, we determined the coefficient of variation (CV) by calculating the variance for each dilution point using log2 data, and then using the average across all dilutions within this range for ranking the pairs. As shown in the annotation table (Supplementary Table 2) the average CV using log-transformed data was 2.2% for plasma and 2% for protein standard and ranged from 0.4% to 7% in plasma and 0.9% to 4.8% in protein. A set of 10 Ab pairs showing averaged CVs for > 3.3% in protein assays and > 4.3% in plasma assays were excluded from further analysis. Prior to choosing the final set of Ab pairs for plasma profiling, sample and protein consumption was considered. Plasma assays requiring more samples > 12.5 μl per assay (representing a 1:4 sample dilution) and amounts of proteins exceeding 150 ng (= 3000 ng/ml as the highest concentration point) were deprioritized. This led to 32 Ab pairs against 22 proteins for further plasma analysis.

#### Preparation of Ab pairs for duplexed plasma analysis

To achieve a more efficient sample analysis, the data from each protein and plasma dilution curve were compared. The concentration levels for a 50% effective dose (ED50) were calculated and chosen as the optimal sample dilution point. In order to find the optimal plasma dilution factor per target protein, one Ab pair and ED50 had to be chosen per protein. In cases of similar performance assessment characteristics (see above), Ab pairs generated towards different binding regions were prioritized. Also, Ab pairs with the superior lower limit of quantification (LLOQ) were preferred as these generally allow us to cover a broader range of protein levels. To attempt for a higher protein throughput, improve time- and cost-efficiency of the assays and also reduce sample consumption, we searched for possible combinations of different Ab pairs with the same optimal sample dilution and limited us to assays in duplex. Some combinations were directly excluded due to previously observed incompatibility, so that 4 duplex assays and 14 single-plex assays remained, as shown in in Table 3. We did not find a direct relation between the protein concentration values found in the literature and the degree of sample dilution (Supplementary Figure 4).

**Table 3.** Determination of secreted proteins in plasma (n = 72).

| Protein | LOD<br>(ng/ml) | LLOQ<br>(ng/ml) | ULOQ<br>(ng/ml) | ED50<br>(ng/ml) | Sample<br>dilution | Determined<br>[c] (ng/ml) | Literature<br>average<br>[c] (ng/ml) | commercial<br>SIA kit range<br>[c] (ng/ml) |
| --- | --- | --- | --- | --- | --- | --- | --- | --- |
| **AFM** | 0.2 | 1.2 | 1,400 | 89.1 | 1/400 | 23,740 ± 7,110 | 75,350[34] | 0.078-5 |
| **ANGPTL3<sup>§</sup>** | 0.01 | 0.05 | 4 | 0.5 | 1/300 | 149 ± 65 | 723[14] | 0.031-2 |
| **APOA4** | 0.06 | 1.1 | 740 | 173.2 | 1/3000 | 32,960 ± 23,170 | 197,500[34] | 0.25-16 |
| **CCL16<sup>§</sup>** | 0.1 | 0.4 | 16 | 2.1 | 1/30 | 71 ± 23 | 11[17] | 0.008-0.5 |
| **CD5L<sup>\*</sup>** | 0.006 | 0.19 | 100 | 18.6 | 1/30 | 441 ± 132 | 5,530[35] | 0.156-10 |
| **CFHR5<sup>\*§</sup>** | 0.1 | 0.4 | 73 | 7.1 | 1/20 | 160 ± 37 | 1660[36] | 0.156-10 |
| **CFP** | 1.3 | 14 | 1,000 | 234.7 | 1/10 | 1,520 ± 467 | 25,000[37] | 15.63-1,000 |
| **CHIT1<sup>\*§</sup>** | 0.002 | 0.01 | 39 | 1.2 | 1/20 | 65 ± 69 | 21.4[11] | 0.156-10 |
| **CPA1<sup>§</sup>** | 0.00003 | 0.008 | 16 | 0.7 | 1/30 | 47 ± 24 | 908.5[4] | 0.65-150 |
| **CPN2** | 0.5 | 2.7 | 910 | 63.1 | 1/1000 | 51,840 ± 13,920 | 55,915.4[4] | 1.56-100 |
| **EFEMP1** | 6.6 | 28 | 1,100 | 97.2 | 1/5 | 381 ± 107 | 10.5[38] | 1.56-100 |
| **EFNA1** | 0.004 | 0.01 | 1.2 | 0.2 | 1/40 | 7.8 ± 2.4 | 7.7[4] | 0.156-10 |
| **FGF21** | 0.00002 | 0.0002 | 0.4 | 0.1 | 1/10 | 0.3 ± 0.2 | 0.5[10] | 0.031-2 |
| **FGL1** | 0.06 | 0.2 | 910 | 47.3 | 1/10 | 613 ± 216 | 23.3[4] | 0.156-10 |
| **GNAS** | 0.003 | 0.01 | 3.6 | 0.7 | 1/10 | 2 ± 0.9 | 5.4[4] | 0.312-20 |
| **HRG** | 0.3 | 1.1 | 23 | 6.7 | 1/1000 | 15,120 ± 2,070 | 100,000[39] | 0.469-30 |
| **HSP90B1<sup>§</sup>** | 0.03 | 0.1 | 4.4 | 1.4 | 1/300 | 972 ± 728 | 464[4] | 0.156-10 |
| **IGFBP2** | 0.02 | 0.2 | 800 | 6.3 | 1/40 | 379 ± 276 | 310[17] | 0.063-4 |
| **INHBC** | 0.02 | 0.1 | 1,100 | 79.0 | 1/40 | 386 ± 194 | 56.2[4] | 0.016-1 |
| **NPPA** | 0.003 | 0.008 | 24 | 0.4 | 1/10 | 9.1 ± 4.5 | 0.06[15] | 0.016-1 |
| **SPON2<sup>\*§</sup>** | 0.2 | 0.6 | 48 | 5.5 | 1/20 | 68 ± 25 | 23.6[40] | 0.078-5 |

(\* measured in serum, <sup>§</sup> measured in duplex assays)

#### Analysis of protein levels in a longitudinal sample set

Lastly, the selected 22 Ab pairs were used in SIAs for the determination of protein levels in a collection of longitudinal plasma samples. The study set was built of 18 individuals that donated plasma every third month over one year. Using the 72 samples collected from four visits each subject allowed us on the one hand to determine the technical suitability of the selected Ab pairs for analysis of proteins, on the other hand we could illustrate how protein levels of individuals vary longitudinally and between sample collections.

We quantified 21 of the tested 22 proteins and listed the performance of the assays in Table 3, where the stated protein concentration for each target was calculated from the average concentration over all samples per donor. The protein levels determined here generally agreed well with those found in the literature (see Supplementary Figure 5). The standard curves from the new assays are shown in Figure 5A-D and relate to those introduced in Figure 4A-D.

**Figure 5A-D.**
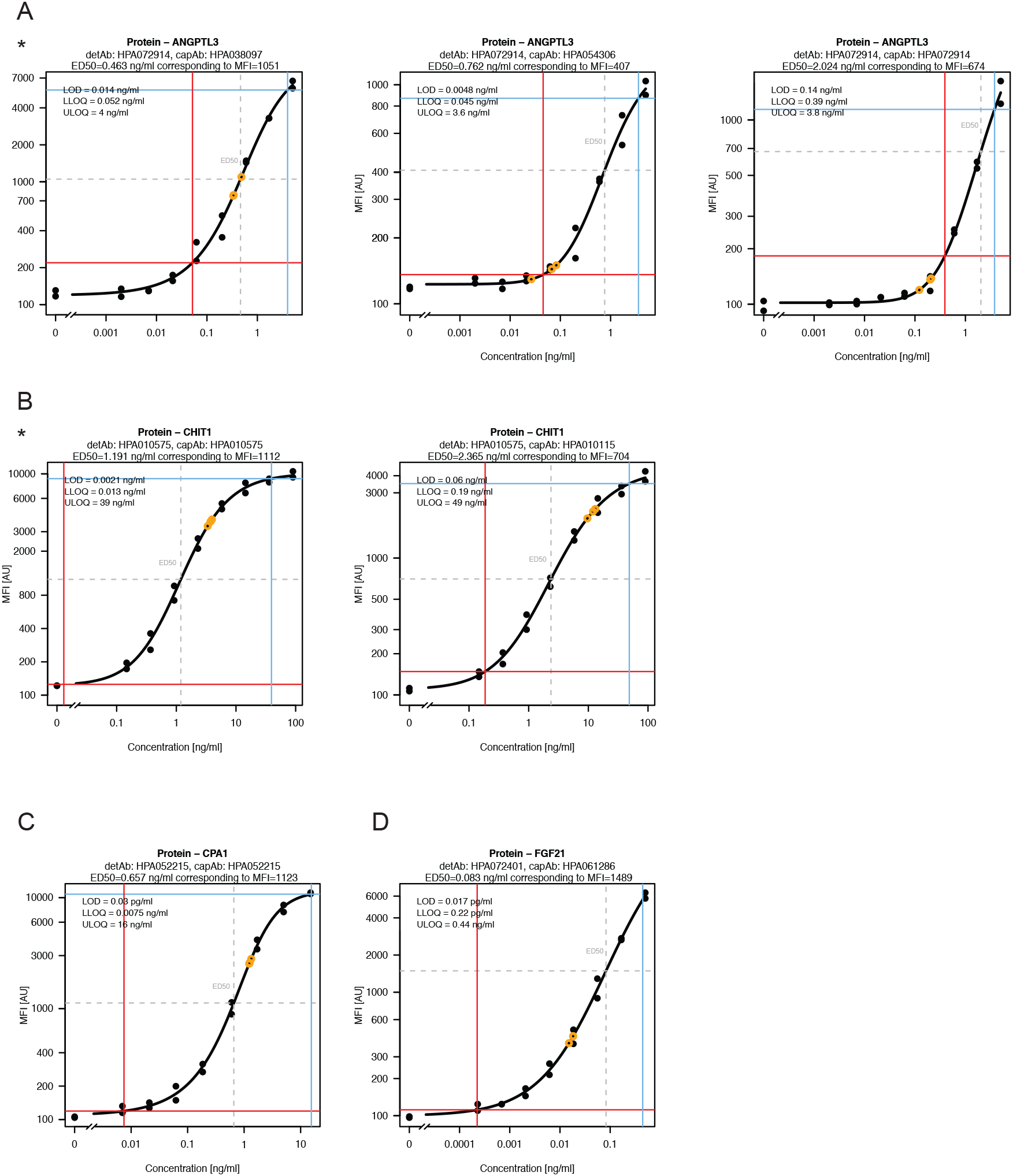
Protein concentration for ANGPTL3, CHIT1, CPA1 and FGF21 in application phase (n = 72). To quantify protein concentration in longitudinal samples during application phase, a 5-parametric log-logistic model was applied for the dilution curves of the protein standard for ANGPTL3 (A), CHIT1 (B), CPA1 (C) and FGF21 (D). Additionally, LOD, LLOQ (red dashed lines), ULOQ (blue dashed lines) and ED50 (grey dashed lines) were calculated. Pooled samples (orange) were plotted onto the curve. If several capAbs were included in the assays, the selected pair is highlighted with *.

In Figure 6 we further compared three layers of variance: technical precision, inter-individual differences as well as longitudinal changes. The ternary plot showed that several proteins, such as CHIT1, CPA1 or FGF21, were stable over time and could be accurately measured, while differing in levels between the donors. The data for ANGPTL3 however was less conclusive due to an elevated technical variance (CV > 21%, using raw data).

In addition to variance analysis, we used distances from clustering analysis to compare the inter-individual differences and the intra-individual differences. Our analysis reveals an average intra-individual Euclidean distance of 3.4 compared to 5.4 between the individuals. This is in line with other observations that protein levels in plasma remain constant over the course of one year and that each person has a unique profile.

**Figure 6.**
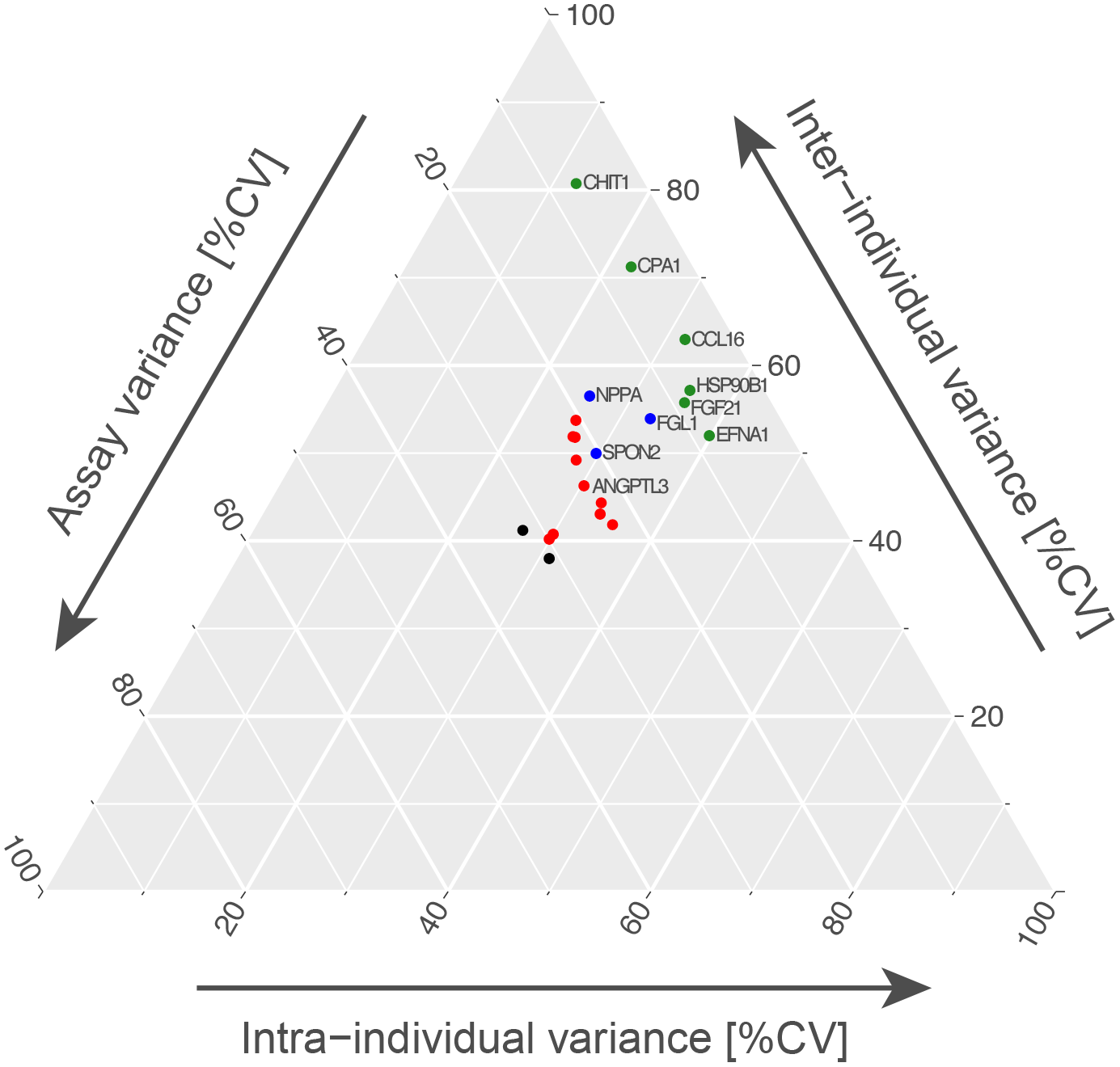
Ternary plot to visualize assay and sample variance. Assay variance was correlated with the inter-individual variance as well as the intra-individual variance using a ternary plot. Assay variance was calculated between the triplicated sample pool, inter-individual CV was defined as the variance of each protein between 18 individuals per visit, while the intra-individual CV is the average variance between the 18 subjects per target over the course of one year. Proteins showing a low assay variance were highlighted in green (0-10%) and blue (10-20%). Data for ANGPTL3 showed an elevated technical variance, which places this protein among ones highlighted in red (assay variance between 20% - 30%).

#### Comparison to orthogonal plasma assays

Lastly, we aimed to confirm the data obtained by the selected Ab pairs though using additional analyses. This assessment was based on comparing our data with results from targeted plasma MS analysis[8] and solution-based proximity extension assays (PEA)[9]. For above methods, we obtained data sets generated in previous studies of the longitudinal sample analyzed in the application phase (Fagerberg et al, unpublished). Using direct correlation analysis as a proxy to determine the similarity between the generated data sets, protein levels from 14 targets were studied. Of the alternative methods, data for 10 proteins only was available for PEA and for 4 proteins from MS only. As shown in Table 4 and Figure 7A-D correlations between our protein levels and another affinity-based method, PEA, reached R^2^ = 0.6 ± 0.2 while correlations with peptide abundance from MS were R^2^ = 0.3 ± 0.2. This illustrates that it was possible to obtain supportive evidence for some of the target proteins, but differences between the assay types in terms of sensitivity and assay interference between the technologies may have contributed to a reduction in concordance. It is worth noting that the assays were performed in different labs and at different timepoints, too. When choosing other capAbs of the SBAs than those shortlisted for the preferred pairs, an additional set of 10 capAbs were available to compare the data from the primary Ab pairs with. Since the data from these assays was obtained from the same sample incubation and used the same detAb, it was less surprising but reassuring to find a high correlation between the primary and additional Ab pairs of R^2^ = 0.9 ± 0.1 (Table 4).

**Figure 7A-D.**
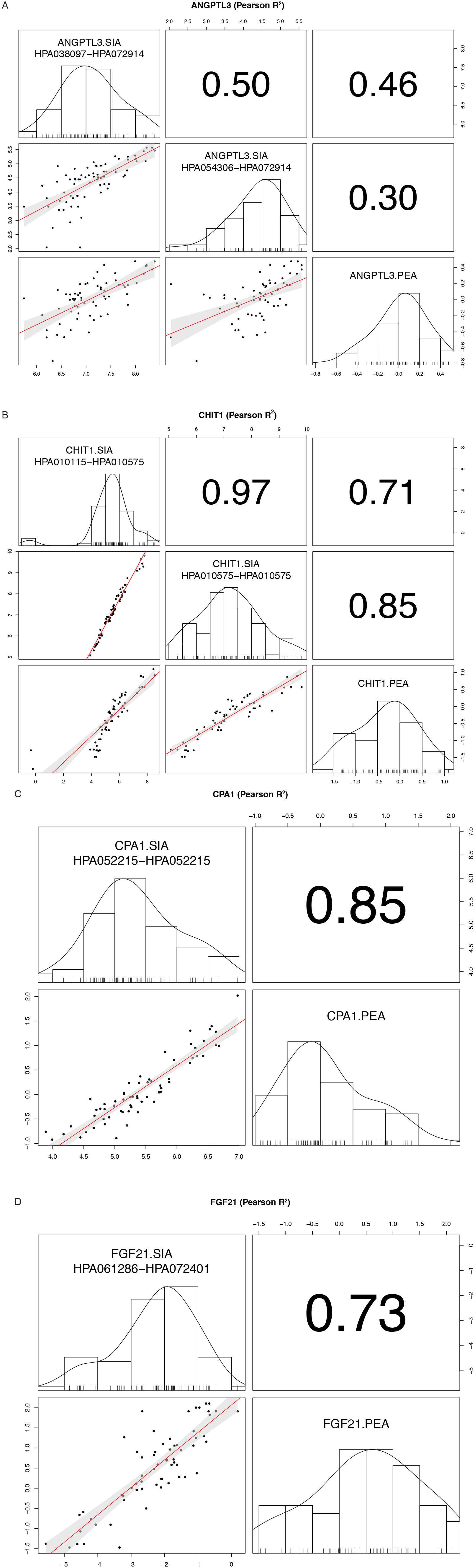
Assay comparison for ANGPTL3, CHIT1, CPA1 and FGF21. Protein concentration achieved from the SIA pairs was compared with data from targeted plasma mass spectrometry analysis and solution-based proximity extension assays, which was generated on the same sample set. 14 out of 21 targets were possible to compare to either of the alternative methods: 4 targets with MS and 10 with PEA. Here, as an example, the correlation for the previously shown 4 targets between SIA and PEA is shown. For visualization and to calculate Pearson correlation R^2^ values, MS data was mean-centered and log2-transformed, SIA data was log2-transformed and NPX values from PEA were used.

**Table 4.**
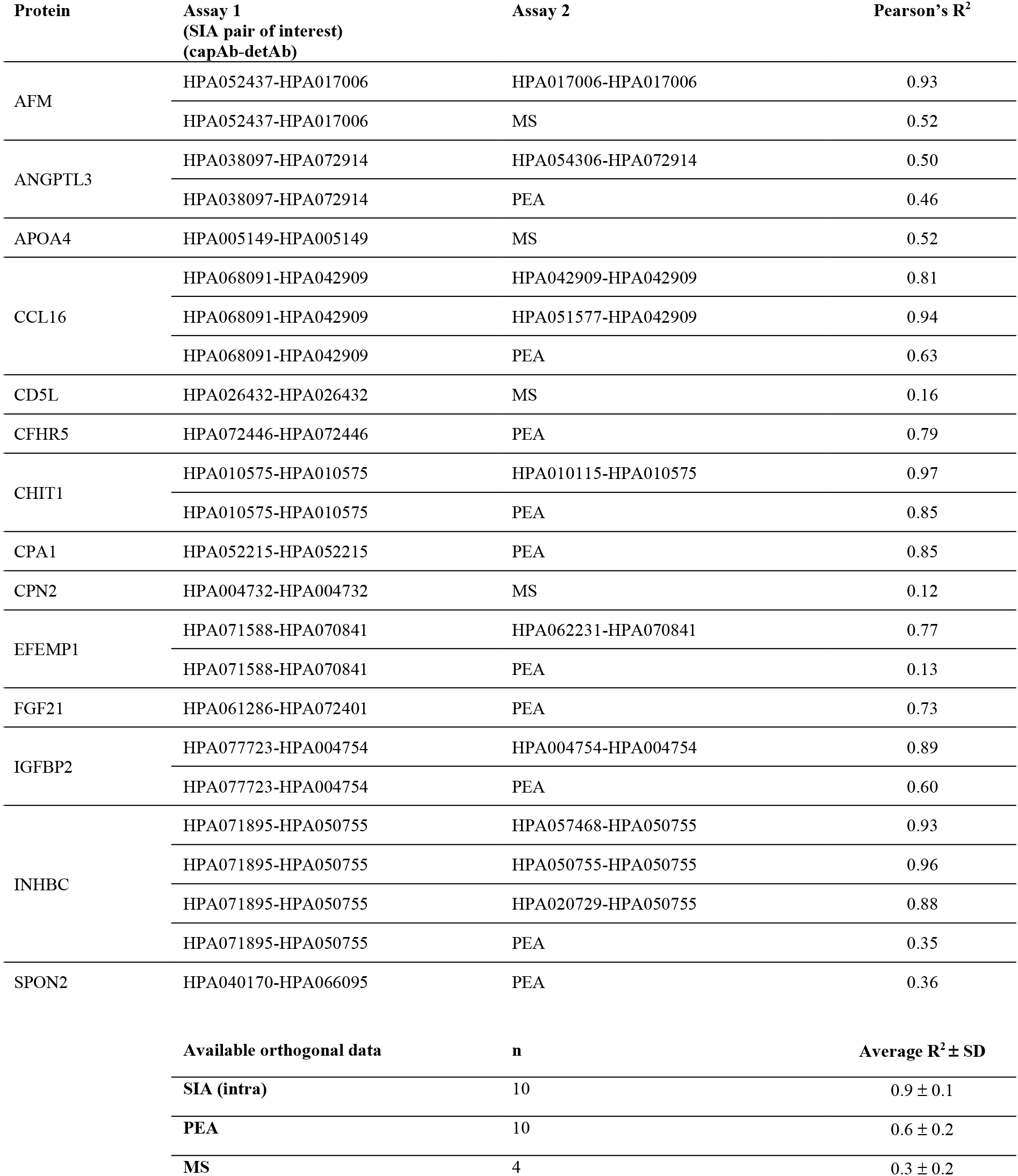
Validation of antibody pair with orthogonal methods (PEA, MS).

## Discussion

This study describes a workflow based on a multiplexed bead-based platform and high-quality reagent resources to systematically screen, select and apply pairs of antibodies for the quantification of secreted proteins in human plasma. Starting from 1,791 unique antibody pairs built on 624 unique antibodies, two rounds of screening were conducted on parallel dilution series of EDTA plasma and the recombinant full-length target protein. We found that 20% or 361 of all possible pairs detected the recombinant as well as the plasma protein in a concentration dependent manner. We applied a selected set of 32 SIA pairs to study the protein levels in plasma collected from 18 subjects every third month over one year and lastly confirmed these findings by using orthogonal assays for 14 targets. For 6 of these, the Pearson correlation between the orthogonal assay and the validated SIA pair was R^2^ > 0.6.

Our study was based on the use of available HSP proteins and HPA antibodies. The polyclonal binders undergo a stringent quality assessment for the use of immunoassays such as Western blot, immunohistochemistry and confocal microscopy. However, the functionality in other types of assays and samples require separate efforts and we did not find a direct link between building functional SIA pairs with pre-assessing these binders in Western blots on plasma (data not shown). Further to this did we not include affinity reagents generated by other providers, which may have limited us in providing larger number of assays in the end. In addition, we estimate that many of the capAbs that are currently part of “non-functional” pairs could indeed enrich the target protein of interest, but the tested detAb was not suitable in combination with these. We also acknowledge the fact that polyclonal antibodies need to be regenerated for extended use and are therefore less suitable for clinical utility. However, applying stringent validation criteria and generation of larger batches may still open up these binders for large scale studies and exploratory research. In addition, identifying suitable antigens from studies based on polyclonal antibodies may streamline the development of monoclonal and recombinant binder libraries.

The proteins that we quantified here are important indicators of health status relevant for different diseases: FGF21 for example is a known key regulator in lipid and glucose metabolism, which is increased in conditions such as type 2 diabetes, obesity and nonalcoholic fatty liver disease[10]. CHIT1 serves as a neuroinflammatory marker and shows increased concentrations in sALS (sporadic amyotrophic lateral sclerosis)[11]. ER stress is hypothesized to lead to hereditary pancreatitis may promote the development of pancreatic ductal adenocarcinoma. *CPA1* is among the highest expressed genes in acinar cells, thus CPA1 proteins are expected to cause more ER stress than lower expressed pancreatic enzymes and are indicated to be associated not only with pancreatic cancer development, but also its susceptibility[12]. ANGPTL3 is a novel factor modulating the plasma lipoprotein metabolism[13] and may additionally contribute to uremic dyslipidemia[14]. Plasma levels of NPPA have been described as prognostic predictors in patients with chronic heart failure, but are also known to reflect the severity of left ventricular hemodynamic dysfunction[15]. It has been suggested that FGL1 plays a role in liver protection and liver regeneration, but it also has the potential to serve as a target for the treatment of gastric cancer and to predict gastric cancer prognosis[16]. In addition, CCL16, a human CC chemokine, has been shown to be differentially expressed in ovarian cancer[17, 18].

The study presented here uses pooled plasma collected from non-diseased subjects. This sample source may have limited the possibility to detect those proteins that increase with inflammation, infection or other diseases. In addition, more assay optimization may have been necessary to rescue some of those pairs that detected their target protein but did not reveal signals above background in plasma. It may further be necessary to choose other, more sensitive detection systems and thereby sacrificing some of the SBA’s capabilities in terms of target and sample throughput. Some Abs showed binding to other than their intended targets once higher amounts of plasma (in particular 1:2). This points at further optimization of the assays are needed in terms of blocking agents and that not all Ab pairs are compatible with another. Additionally, the apparent LOD and LLOQ levels may increase once more complex buffer solutions are used.

To our current knowledge, this is one of the largest and first systematic study to screen for SIAs. We focused on the plasma secretome, as plasma is an important sample for clinical routine analysis and a highly interesting source for searching for disease related proteins. We assumed that the proteins secreted into the blood stream remain detectable in solution when plasma is being prepared. In comparison to studying proteins in cell lysates, proteins found in plasma do not require to be extracted, hence the need to apply strong detergents can be omitted. However, we acknowledge that proteins may precipitate or denature during sample processing. Some proteins are known to be unstable and degrade over time, hence the possibility to detect these decreases with the age of the sample.

In addition to expand our possibilities to measure proteins actively secreted into blood, highly multiplexed immunoassays as well as MS-based approaches require more targeted assays to validate and quantify potential findings in larger number of samples as well as using orthogonal methods. It holds a great value to expand this list of assays for the plasma secretome to measure and quantify the plasma components of interest on a protein level. Considering the targets included in our study, this suggests that the detectability of proteins in plasma is predominantly depended on the interplay between the available reagents, the technology and protein stability. We are, however, well aware that many other, such as the low abundant cytokines[19], require minimal sample dilutions to detect proteins of pg/ml concentrations, and we suggest to include necessary controls and considerate elevated and unspecific background binding.

In summary, multiplexed bead arrays were used to screen for functional antibody pairs in proteins and plasma samples. With a success rate of 20% we found that investing at least three different antibodies per target protein and assessing different capture-detection combinations was necessary to obtain antibody pairs for protein quantification. While further assay optimization, additional antibodies and target-centric studies will be needed to assess the utility of these antibody pairs, we could not observe a trend towards any binding-site preference for building functional assays for secreted proteins. Considering the need to generate renewable reagents for extended use, our study provides valuable leads on selecting and building antibody pairs even with other reagents than those used here.

## Methods

### Plasma samples

All methods were conducted according to the Declaration of Helsinki, which establishes the regulations and guidelines for research project execution for human health.

The screening for SIA pairs was conducted on pools of anonymous donors and did not require sensitive personal information about the donors. The research did not include any type of intervention, surgery or treatment. The Ethical Review Board in Uppsala (Dnr 2009/019) deemed that this research was not subjected to formal ethical review and approval. Samples of human K2 EDTA plasma were purchased on two occasions from Sera Laboratories International Ltd (HMPLEDTA2, now part of BioIVT, West Sussex, UK), who collects samples under IRB-approved protocols in use at their FDA-licensed donor centers with written informed consent obtained from all donors. The pools of plasma samples were generated by the supplier from mixing plasma from donors of which 50% were females.

The selected SIA pairs were then used to study samples collected from 18 subjects over a one-year time period. The SIA pair targeting EFEMP1 was run on a different selection of 18 subjects, due to the available sample volume. Each subject donated plasma every third month as part of the longitudinal Swedish SCAPIS SciLifeLab Wellness Profiling (S3WP) program (Dnr 407-15). Within this study a total of 101 subjects were recruited from the ongoing Swedish CArdioPulmonary bioImage Study (SCAPIS), which is a prospective observational study of randomly selected subjects aged 50-64 years from the general Swedish population. Within this study, all subjects have been extensively phenotyped before entering the S3WP program[20]. Several exclusion criteria were applied for choosing study participants: 1) previously received health care for myocardial infarction, stroke, peripheral artery disease or diabetes, 2) presence of any clinically significant disease which, in the opinion of the investigator, may interfere with the results or the subject’s ability to participate in the study, 3) any major surgical procedure or trauma within 4 weeks of the first study visit, or 4) medication for hypertension or hyperlipidemia. The study was approved by the Ethical Review Board of Göteborg, Sweden. All participants provided written informed consent. The S3WP program has the aim to collect longitudinal data in a community-based cohort and is non-interventional and observational. A total of 4 examinations were performed every third month (+/- 2 weeks). All subjects were fasting overnight (at least 8 hours) before the visits. Subjects underwent the same examinations at each visit, answered a selection of questions to note changes in health and life-style factors between each visit and blood, urine and stool for subsequent clinical chemistry and omics analyses was collected at each visit (Fagerberg et al, unpublished).

### Target selection and generation

Protein targets for the secretome were selected according to availability of full-length proteins within the Human Secretome Project (HSP) and antibodies from the Human Protein Atlas (HPA) as well as considering the recombinant protein concentration. HPA antibodies needed to have a concentration of > 0.05 mg/ml for being chosen as capAb and of > 0.1 mg/ml for being considered as detAb.

### Production and purification of secreted proteins

Secreted proteins were defined based on data in the Uniprot database as well as signal peptide and transmembrane region predictions made for the transcripts in the Ensembl database. A generic expression cassette, based on the CMV promoter and with an N-terminal CD33 signal peptide for secretion of all produced proteins and a C-terminal human protein C tag for purification, was used. All secreted proteins were produced by using the transient Icosagen Cell Factory system with CHOEBNALT-85 cells and the QMCF Technology (Icosagen Cell Factory OÜ, Tartu, Estonia). Cells were maintained in a 50:50 mixture of 293 SFM II (Gibco, 11686029) and CD CHO medium (Gibco, 10743001) with a supplement of 6 mM GlutaMAX (Gibco, 35050061) and 10 ml/l HT supplement 50X (Gibco, 41065012) at 37°C on an orbital shaker. A total of 6 million cells were transfected by electroporation. The transfected cells were added to fresh pre-warmed 20 ml medium containing penicillin-streptomycin (Sigma Aldrich, P4333-100ML) in 125 ml shaking flasks (Sigma Aldrich, CLS431143-50EA) and cultivated in a fed batch cultivation for 13 days. 48 h after transfection cells were diluted to 400,000 cells/ml with fresh medium. Successful transfection and protein secretion were determined six days after transfection by performing Western Blots. Positive screened samples were initiated to production by the addition of 20% CHO CD EfficientFeed B (Thermo Fisher, A1024001) and a temperature shift to 30°C. A second feed of 10% was added at day 9 after transfection. The supernatant was clarified by centrifugation and serine-protease inhibitor was then added. For purification 1 ml of an in house developed anti protein C affinity matrix was used. The harvest sample was filtrated into the matrix and CaCl_2_ was added to a final concentration of 2 mM. The tube with sample and matrix was then incubated in a cold room overnight. After packing the matrix in a column, it was washed with equilibration buffer (20 mM Tris, 100 mM NaCl, 2 mM CaCl2, pH 7.5) and thereafter a filter was placed on top of the matrix and the column was placed on ASPEC 271 or 274 liquid handlers (Gilson Inc.). After an additional washing step (20 mM Tris, 1 M NaCl, 2 mM CaCl_2_, pH 7.5) the protein was eluted using a mild elution with EDTA (20 mM Tris, 100 mM NaCl, 2 mM EDTA, pH 7.5) prior a buffer exchange into 1xPBS. After desalting the protein concentration was determined (Abs). Each purified protein was identified by MS/MS and the purity was analyzed using SDS-PAGE and Western Blot. Primary antibody for western blotting was a rabbit Anti-C tag polyclonal (GTX18591, Genetex). Glycosylation patterns of the purified proteins were also analysed using SDS-PAGE.

### Antibodies

Overall, we included 624 antibodies targeting 209 unique secreted human proteins, as well as 11 assay specific controls. Majority of the antibodies used polyclonal rabbit antibodies generated within Human Protein Atlas project (HPA) (www.proteinatlas.org)[21]. The assay specific controls included affinity purified rabbit IgG (P120-301, Bethyl laboratories) in order to control for background binding to rabbit IgG molecules and a blocked bare bead (without coupled antibody) to monitor background binding to the beads. These two will from now on be referred to as “assay controls”. In addition, a set of 10 monoclonal mouse antibodies from BioSystems International[22], targeting plasma proteins commonly enriched by immuno-capture assays were included[23]. These will be referred to as “internal controls”. All antibodies used are listed in (Supplementary Table 1).

### Coupling of antibodies to beads

Bead arrays were created as previously described[24]. Antibodies were diluted to 17.5 μg/ml in 100 μl 0.1M 2[N-Morpholino]ethanesulfonic acid (MES)-buffer (M2933, Sigma-Aldrich), pH 4.5, using a pipetting robot (TECAN EVO150) and then coupled to carboxylated color-coded magnetic beads (MagPlex-C, Luminex Corp). In short, beads (n = 500 000/ID) located in 96-well microtiter plates (Greiner BioOne) were washed with 80 μl 0.1M NaH2PO4 (phosphate buffer) pH 6.2 (S3139, Sigma Life Science) with a plate washer/dispenser (EL406, Biotek) on magnet. Subsequently, 50 μl phosphate-buffer was added manually. Activation buffer consisting of 10 mg/ml 1-ethyl-3-(3-dimethylaminopropyl) carbodiimide (EDC) (C1100, ProteoChem) and 10 mg/ml Sulfo-N-hydroxysulfosuccinimide (Sulfo-NHS) (24510, Thermo-Fisher Scientific) in phosphate buffer were subsequently added to the beads, resulting in 0.5 mg EDC and 0.5 mg Sulfo-NHS per well. Activation buffer and beads were incubated for 20 min at 650 rpm at room temperature and washed two times with 100 μl 0.1M MES. The pre-diluted antibodies were added to the activated beads and incubated for 2 h at 650 rpm at room temperature. After incubation, the antibody-coupled beads were washed three times in 100 μl 1x PBS (09-9400, Medicago), 0.05% Tween20 (BP337, Fisher Bioreagents) (PBS-T) and re-suspended in 50 μl storage buffer (Blocking Reagent for ELISA, 11 112 589 001, Roche Diagnostics) supplemented with ProClin (4812-U, Sigma-Aldrich). The individual bead IDs were pooled together after overnight blocking at 4°C, creating six bead stocks containing 65-95 different kinds of antibody coupled beads, including 10 additional control antibodies each, coupled to unique bead IDs. The coupling efficiency of the antibody-coupled beads was tested using R-Phycoerythrin-conjugated (RPE) goat anti-rabbit IgG (111-116-144, Jackson ImmunoResearch) and RPE-conjugated goat anti-mouse IgG (115-116-146, Jackson ImmunoResearch). 100 μl RPE-conjugated antibodies diluted to 0.5 μg/ml in PBS-T were added to 5 μl antibody-coupled bead stock in different wells, followed by incubation for 20 min at 650 rpm at room temperature. After incubation, wells were washed three times with 100 μl PBS-T before analyzed on a Flexmap 3D instrument (Luminex corp.). Signals for the coupling efficiency was reported in terms of median fluorescence intensities (MFI). Coupled beads were regarded as a failed coupling if the signals obtained were lower than 2x SD than the mean value for the bead stock. In case of failed coupling, a recoupling of this specific antibody was performed.

### Biotinylation of detection antibodies

Antibodies used as detAbs were biotinylated as described previously[25]. In short, 2 μg of each antibody was diluted in 30 μl PBS-T and then incubated with 5 μl protein A-coated magnetic beads (30 mg/ml, Dynabeads, 10002D, Invitrogen) for 30 min, room temperature, 650 rpm. After incubation, the antibody-coupled beads were washed three times in 100 μl PBS-T before labelling the antibodies with a 150x molar excess of EZ-Link-NHS-PEG4-Biotin (21329, Thermo Scientific) dissolved in DMSO (276855, Sigma-Aldrich) for 30 min, room temperature, 650 rpm. The beads were then washed three times in 100 μl PBS-T. The labelled antibodies were dissociated from the beads by adding 15 μl 0.2M acetate (97064-482, VWR), pH 3.2 (elution buffer) to the wells and incubated for 2 mins at room temperature, while mixing gently. The supernatants were collected using a magnet and transferred into individual tubes. To buffer the solution 5 μl of 0.5M Tris-base (T6066, Sigma-Aldrich), pH 8 were added to each eluate. Subsequently, 5 μl PBS-T were added to each tube and the labelled antibodies were stored at 4°C with an estimated concentration of 0.072 μg/μl.

The biotinylation efficiency was tested by diluting the labelled antibody to 1 μg/ml in PBS-T, and adding 25 μl of pre-diluted labelled antibody to 2 μl of donkey anti-rabbit IgG (711-005-152, Jackson ImmunoResearch) coupled beads followed by incubation for 1 h at 650 rpm at room temperature. After incubation, wells were washed three times with 100 μl PBS-T.

Subsequently, 50 μl of a 1:750 dilution of RPE-labeled streptavidin (SA10044, Invitrogen) were added and incubated for 20 min, room temperature, 650rpm. After incubation, wells were washed three times with 100 μl PBS-T, before analyzed on a Flexmap 3D instrument (Luminex corp.). Antibodies with signals 50 x above background were considered successfully biotinylated.

### Assay procedure and read out

For assay performance two different batches of commercially available human K2 EDTA mixed gender plasma pool (HMPLEDTA2, Seralab) were serially diluted in plasma dilution buffer to cover a dilution range of 1:4 till 1:3000 in seven steps with equal dilution. The first screening round used a different plasma batch then the rest of the experiments. The plasma dilution buffer consisted of 1x PBS with 0.5% (w/v) polyvinylalcohol (P8136, Sigma-Aldrich), 0.8% (w/v) polyvinylpyrrolidone (PVP360, Sigma Life Science), 0.1% casein (C5890, Sigma Life Science) and supplemented with 0.5 mg/ml rabbit IgG. A spike-in serial dilution of standard proteins in plasma dilution buffer was performed, covering a concentration range of 1 μg/ml − 1 ng/ml. Blanks of both assay buffer and plasma dilution buffer were added and will be referred to as “blank sample”. The pre-diluted plasma samples and the pre-diluted protein standards (45 μl) were transferred to 5 μl bead stock in an assay plate (Greiner 384-well assay plate) using a liquid handler (SELMA, CyBio) before overnight incubation at 650 rpm at room temperature.

After incubation, the beads were washed three times with 60 μl PBS-T. The biotinylated detAbs were diluted to 1 μg/ml in PBS-T. Subsequently the beads were incubated for 1.5 h at room temperature at 650 rpm with 25 μl of the respective pre-diluted detAb. Beads were washed three times with 60 μl PBS-T before incubation with 50 μl of a 1:750 dilution of RPE-labelled streptavidin for 20 min at room temperature at 650 rpm. Finally, beads were washed three times with 60 μl PBS-T, before they were re-suspended in 60 μl PBS-T and analyzed on a Flexmap 3D instrument (Luminex corp.). Binding events were displayed as MFI where at least 50 beads per bead ID were counted.

Each assay plate represents one experimental assay run combined of 2x 96-well plates containing serial dilutions of human plasma pool as well as 2x 96-well plates containing serial diluted standard curves of the proteins investigated. Protein and plasma dilution series for the same detAbs were placed on the same 384-plate (Supplementary Figure 1). Additionally, interfering plate-effects were avoided by running all measurements for one target protein on the same 384-plate. In total, 26 384-well plates were measured, containing between 2 and 15 proteins each. A detailed plate layout can be found in the Supplementary Figure 1.

### Assay optimization and validation

The assay design was modified and optimized each time between two phases. After the screening phase, the buffer matrix for the protein standards was changed by adding 1% BSA (A7030, Sigma Life Science) to achieve a higher matrix complexity. Additionally, the length of the dilution curves for the protein standard was extended from a 7-step concentration to a 14-step concentration series in triplicates covering a range of 1 μg/ml to 1 pg/ml when evaluating the reproducibility of the assays. The dilution points for plasma were also adapted to the signals achieved during screening, to both cover a broader measuring range, but also to be more suitable for the obtained signals. Thus a 7-step dilution series of human EDTA plasma pool with a consistent step size of 3 starting between 1:2 and 1:36 in plasma dilution buffer was conducted for each antibody pair. The SBAs for the selection process were composed with different capAbs for further technical investigations in order to exclude additional off-target interactions. The remaining target proteins were grouped as sets of five into SBAs containing 8-18 Abs, based on their alphabetical order. Each set of SBAs was supplemented with the assay controls, to record possible binding to the beads. For protein quantification and application of the SIA pairs, assays were run on a longitudinal sample set as well as an 8-step protein concentration series, covering the optimized signal range and measured in triplicates.

### Selection criteria

After selecting protein targets for the secretome according to availability, and HPA antibodies both according to availability and concentration (> 0.05 mg/ml for being chosen either as capAb or detAb), antibody pairs had passed several selection rounds in order to achieve reliably functioning antibody pairs. This process was divided into three phases: an initial screening phase, which was sub-divided into two rounds, a selection phase and an application phase. For all phases MFIs were registered for each bead ID and sample.

Annotation after screening phase was performed manually. Hereby we grouped antibody pairs based on their functionality into 4 different categories: 1) Dilution dependent curves with protein and plasma, 2) Dilution dependent curves with protein only, 3) Dilution dependent curves with plasma only and 4) No dilution dependent curves. We assessed this according to the shape and concentration dependency of the curve for the expected pair.

Pairs being processed to be further tested had to be assigned to group 1 as well as reach a maximum signal intensity of at least 150 MFI in order to report only signals above an average background. To limit the number of total assays, one detAb was chosen per protein. For the detAbs, any concentration dependent binding for the other Ab-coupled beads in the SBA, such as the internal controls, were used as exclusion criteria. As additional criteria a pair was chosen upon showing the widest range of detectable concentrations of proteins in buffer and plasma, an overall lower background level in antigen-free samples, and indications about possible interferences or off-target recognition of other captured proteins. Lastly, the available antibody volume was considered for the polyclonal binders.

After an additional testing round of the chosen pairs in triplicates of plasma dilution and protein standard dilution the CV was determined for each dilution point averaging it across all dilution steps to find the best performing Ab pairs (see Figure 4A-D). As cut-off criteria a CV of 3.3% in protein assays and 4.3% in plasma assays was defined. In addition, pairs requiring more than 12.5 μl sample (representing a 1:4 sample dilution) or more than 150 ng (= 3000 ng/ml as the highest concentration point) per assay were excluded in respect to sample and protein consumption. For the technical replicates during the selection phase the upper limit of quantification (ULOQ), LLOQ, LOD and ED50 were calculated. One pair per target was prioritized and the calculated ED50 point was chosen as the optimal sample dilution point. In cases of similar performance, Ab pairs generated towards different binding regions were prioritized. Also, Ab pairs with the superior LLOQ were preferred (see Supplementary Table 2).

Before processing the remaining Ab pairs into the final application phase and measuring them on a selection of 72 samples from a healthy longitudinal cohort, we combined different Ab pairs into possible duplex combinations with the same determined optimal sample dilution. The final concentration measured for each sample was calculated by transforming the measured MFI signal intensities into a concentration value according to the dilution curve obtained from the 5-parametic fit and multiplying it with the applied dilution factor of the sample (see Figure 5A-D). Samples with protein concentration below the calculated LLOQ or above ULOQ were excluded from further analysis.

### Orthogonal assays

For orthogonal comparison of our targets, we correlated the overlap of our chosen 21 targets with data achieved by two independent experimental setups for the same sample selection: (1) a recently published targeted MS approach[8] and (2) multiplex proximity extension assays (Olink Bioscience, Uppsala Sweden)[9].

For the targeted MS approach 432 samples were prepared semi-automatically using the Bravo liquid handler and subsequently measured using a combination of Ultimate 3000 binary RS nano liquid chromatography (LC) system (Thermo Scientific) with an EASY-Spray ion source connected to an on-line Q Exactive HF (Thermo Scientific) MS. All plasma samples were stored lyophilized and resuspended by the autosampler. Sample analysis was performed using a previously developed PRM method. Each full MS scan at 60,000 resolution (AGC target 3e^6^, mass range 350-1,600 m/z and injection time 110 ms) was followed by 20 MS/MS scans at 30,0 resolution (AGC target 2e^5^, NCE 27, isolation window 1.5 m/z and injection time 55 ms) which were defined by a scheduled (2 min windows) PRM isolation list that contained 174 paired light and heavy peptide precursors (n(peptides) = 87) from 55 QPrESTs directed towards 52 human proteins. The raw MS-files from all study samples were processed in Skyline (version 3.7) and analyzed in R (version 3.4.1) for protein quantification.

For some of the measured plasma proteins additional validation was achieved by using multiplex proximity extension assays. Each kit contains a microtiter plate measuring 92 protein biomarkers in up to 90 samples. Each well contains 96 pairs of DNA-labeled antibody probes. Samples were incubated in the presence of proximity antibody pairs tagged with DNA reporter molecules. When the antibodies pair binds to their corresponding antigens, the corresponding DNA tails form an amplicon by proximity extension, which can be quantified by high-throughput real-time PCR[9, 26]. To minimize inter-and intra-run variation, the data are normalized using both an internal control (extension control) and an interplate control, and then transformed using a pre-determined correction factor. The pre-processed data were provided in the arbitrary unit Normalized Protein expression (NPX) on a log2 scale. A high NPX presents high protein concentration[26].

### Data analysis (Data processing, Classification and Curve fitting)

Data analysis and visualizations were performed within R (www.rproject.org, version R 3.5.1)[27]. To assess reproducibility for overlapping targets between the two screening rounds, corresponding MFI values were log transformed and correlated using Pearson correlation with R^2^ values. To assess the binding region for capAbs and detAbs on the screened proteins, the immunogens aminoacidic sequence for each HPA was mapped to the sequences of the corresponding canonical protein (www.proteinatlas.org). Protein sequences were exported from the Uniprot data base (release 2018_07)[28].

To evaluate the performance of Ab pairs during the selection phase, data was log10 transformed and visualized as dilution curves.

Data was log10 transformed and a 5-parametric log-logistic model was applied for the dilution curves in the application phase[29]. LOD levels were calculated as 3x SD of the blank sample above the average blank signal, LLOQ was defined as 10x SD of the blank sample above the average blank signal[30], ULOQ was defined as the averaged signal of the highest protein standard concentration point minus its SD. ED50 was calculated using the drc package[29]. In instances where the SD was small, leading to negative output of the 5-parametric fit for the LOD values, the MFI values for the blank were manually increase by adding 2 AU. No significant effect on the calculated protein concentrations for those targets could be observed.

Assay CVs within the selection phase were calculated between the duplicated dilutions steps of the plasma protein curve, while during the later application phase the assay variance was calculated with the triplicated sample pool. During this phase two additional layers of variance were calculated: Variances for each protein between the 18 individuals (per visit), which will be referred to as inter-individual CV, as well as the average variance between the 18 subjects over the course of one year (four samples per subject), which we will refer to as intra-individual variance. For visualizing different layers of variance (assay variance, inter-individual variance and intra-individual variance), CVs were calculated and ternary plots were generated using the ggtern package[31].

Euclidian distances for investigating the personal plasma profile differences were calculated using the daisy function[32, 33]. Prior to the calculation, the data underwent an outlier removal process, meaning values above ULOQ and below LLOQ as well as NA values were removed from the data set. The data was then scaled before computing pairwise dissimilarities and Euclidean distances. Additionally, Pearson distance for inter-individual and intra-individual correlation was calculated, using the R^2^ value.

For the correlation plots between different types of data were used: MS data (fmol/μl) was mean-centered and both MS data as well as SIA data (ng/ml) were log-transformed with the binary logarithm, while PEA data (NPX values) was used as provided and correlated using Pearson correlation with R^2^ values.

## Supporting information

Supplementary Figure 1. Plate layout for screening phase

Supplementary Figure 2. Number of tested pairs per unique target

Supplementary Figure 3. Functional SIA pairs screening # 1 and # 2

Supplementary Figure 4. Correlation between protein concentration published in literature and the degree of sample dilution

Supplementary Figure 5. Correlation between measured protein concentration and the protein concentration published in literature

Supplementary Table 1. All antibodies screened

Supplementary Table 2. Annotation table - pair selection for application phase

## Acknowledgements

We thank everyone at the Affinity Proteomics and Clinically Applied Proteomics groups at SciLifeLab in Stockholm for their continuous fruitful discussion, access to instrumentation and input to the presented work. We also thank everyone at the Human Protein Atlas (HPA) and at the Human Secretome Project (HSP) for their support. The KTH Center for Applied Precision Medicine (KCAP) funded by the Erling-Persson Family Foundation is acknowledged for financial support. This work was supported by grants for Science for Life Laboratory, and the Knut and Alice Wallenberg Foundation for funding the HPA project and the HSP. HSP was also funded by AstraZeneca and Novo Nordisk Foundation. The work leading to this publication has received support from the Innovative Medicines Initiative Joint under grant agreement n°115317 (DIRECT), resources of which are composed of financial contribution from the European Union’s Seventh Framework Programme (FP7/2007-2013) and EFPIA companies’ in-kind contribution.

## Author Contributions Statement

RSH, AB, MJI, LS, MD, EB, SB, CF planned and performed immunoassays. RSH, AB, UQ and JMS analyzed data. FE performed MS assays and analysis. TDC analyzed data. LF provided Olink data. JR and HT supervised the protein production and HT provided the project with full-length proteins. MU and JMS conceived the study. UQ and JMS supervised the study. RSH, AB and JMS wrote the manuscript with scientific input from all co-authors.

## Additional information

UQ is employee of Atlas Antibodies AB. MU is co-founder of Atlas Antibodies AB. FE, LF, JR, HT and JMS acknowledge formal links to Atlas Antibodies AB.

## Abbreviations

Abs: Antibodies
capAb: capture Antibody
CV: Coefficient of Variation
detAb: detection Antibody
HPA: Human Protein Atlas
HSP: Human Secretome Project
MFI: Median Fluorescence Intensity
MS: Mass Spectrometry
PEA: solution-based Proximity Extension Assay
S3WP: Swedish SCAPIS SciLifeLab Wellness Profiling
SBA: Suspension Bead Array
SIA: Sandwich Immunoassay

## Supplementary Figure Legends

**Supplementary Figure 1. Plate layout for screening phase.**

During screening phase 4x 96-well plates were combined into one 384w-plate, representing one experimental run. Each run was combined of 2x 96-well plates containing serial dilutions of human plasma pool as well as 2x 96-well plates containing serial diluted standard curves of the proteins investigated. Protein and plasma dilution series for the same detAbs were placed on the same 384-plate.

**Supplementary Figure 2. Number of tested pairs per unique target.**

Different amounts of Ab pairs were tested for the different targets according to availability (n = 1-100).

**Supplementary Figure 3. Functional SIA pairs screening # 1 and # 2.**

Ab pairs tested both in plasma and towards full-length protein. 0 = pair is not functional, 1 = pair shows a concentration dependent curve and is regarded as functional.

**Supplementary Figure 4. Correlation between protein concentration published in literature and the degree of sample dilution.**

The degree of dilution determined according to ED50 per protein was correlated with the protein concentration according to literature. Axes are log-scaled and a line of identity was added to the plot.

**Supplementary Figure 5. Correlation between measured protein concentration and the protein concentration published in literature.**

The determined protein concentration was correlated to concentrations for the same protein according to literature. Axes are log-scaled and a line of identity was added to the plot.

**Supplementary Table 1. All antibodies screened.**

**Supplementary Table 2. Annotation table - pair selection for application phase.**

Ab pairs showing a signal intensity of MFI > 150 for the protein assay during the screening phase are listed as well as their exclusion criteria during the selection phase. Pairs in red were excluded before application phase, black and green pairs were processed to the subsequent application phase. Green pairs represent the final selected pairs for application phase according to ED50 and LLOQ. Exclusion reason for each antibody is indicated with x.

