## Supplementary figures and images for "Systematic Development of Sandwich Immunoassays for the Plasma Secretome"

### Supplementary Figure 1. Plate layout for screening phase

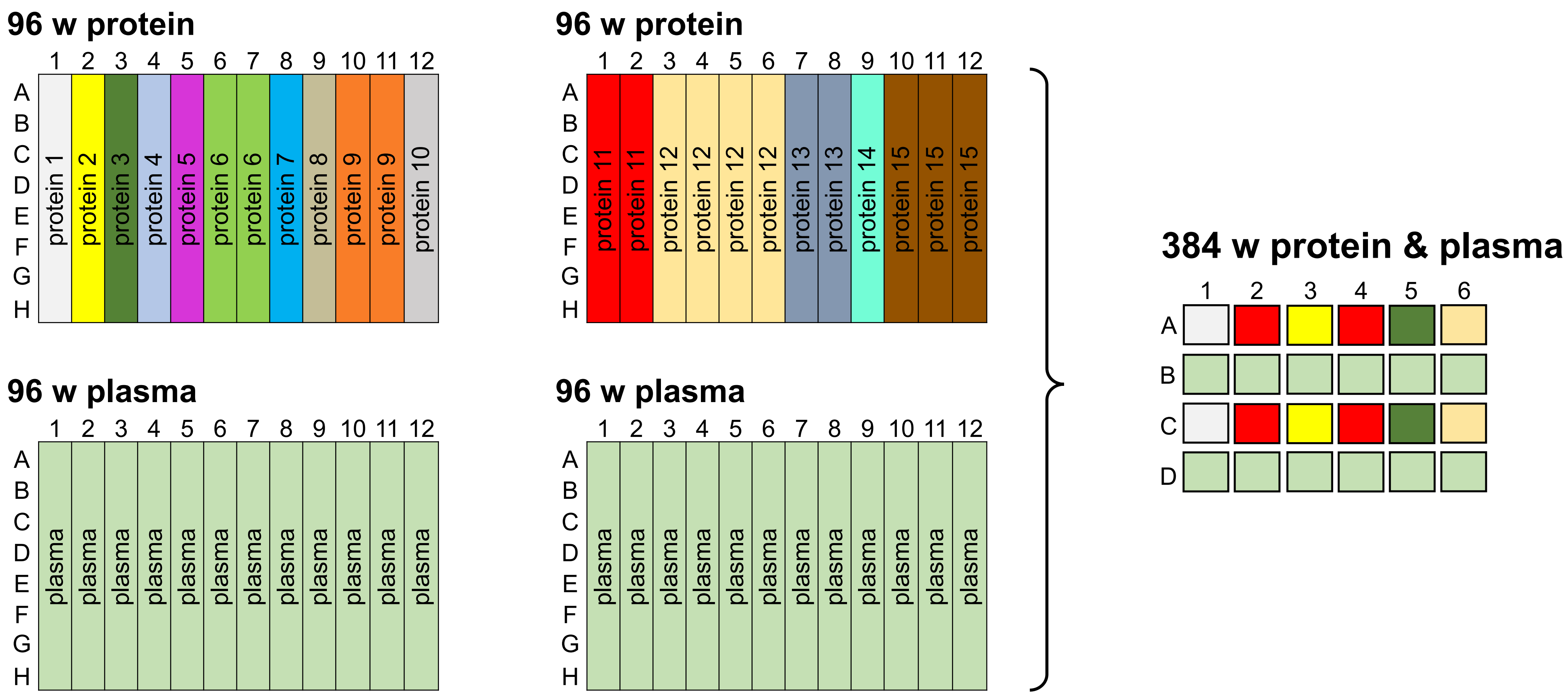

### Supplementary Figure 4. Correlation between protein concentration published in literature and the degree of sample dilution

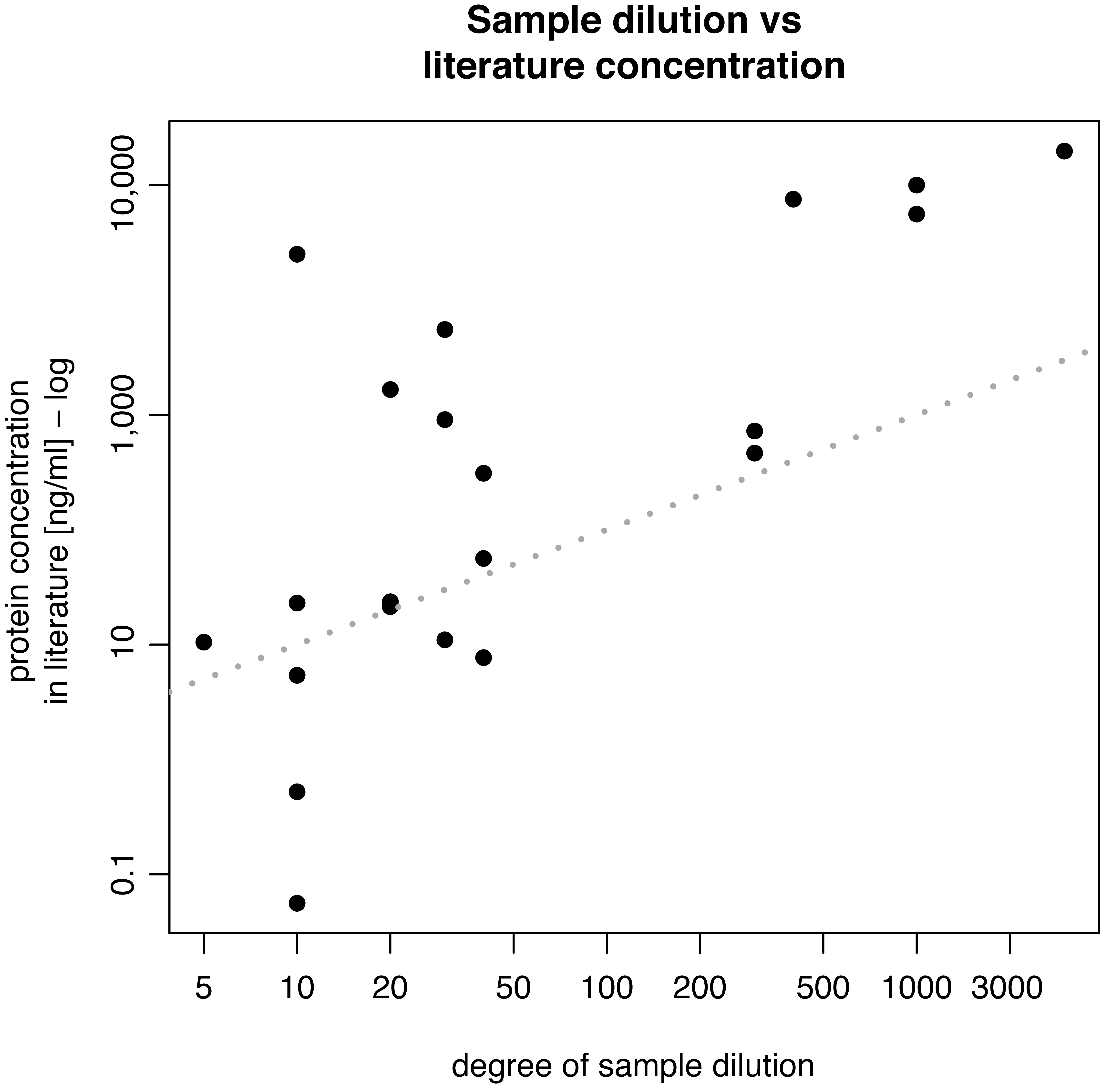

### Supplementary Figure 5. Correlation between measured protein concentration and the protein concentration published in literature

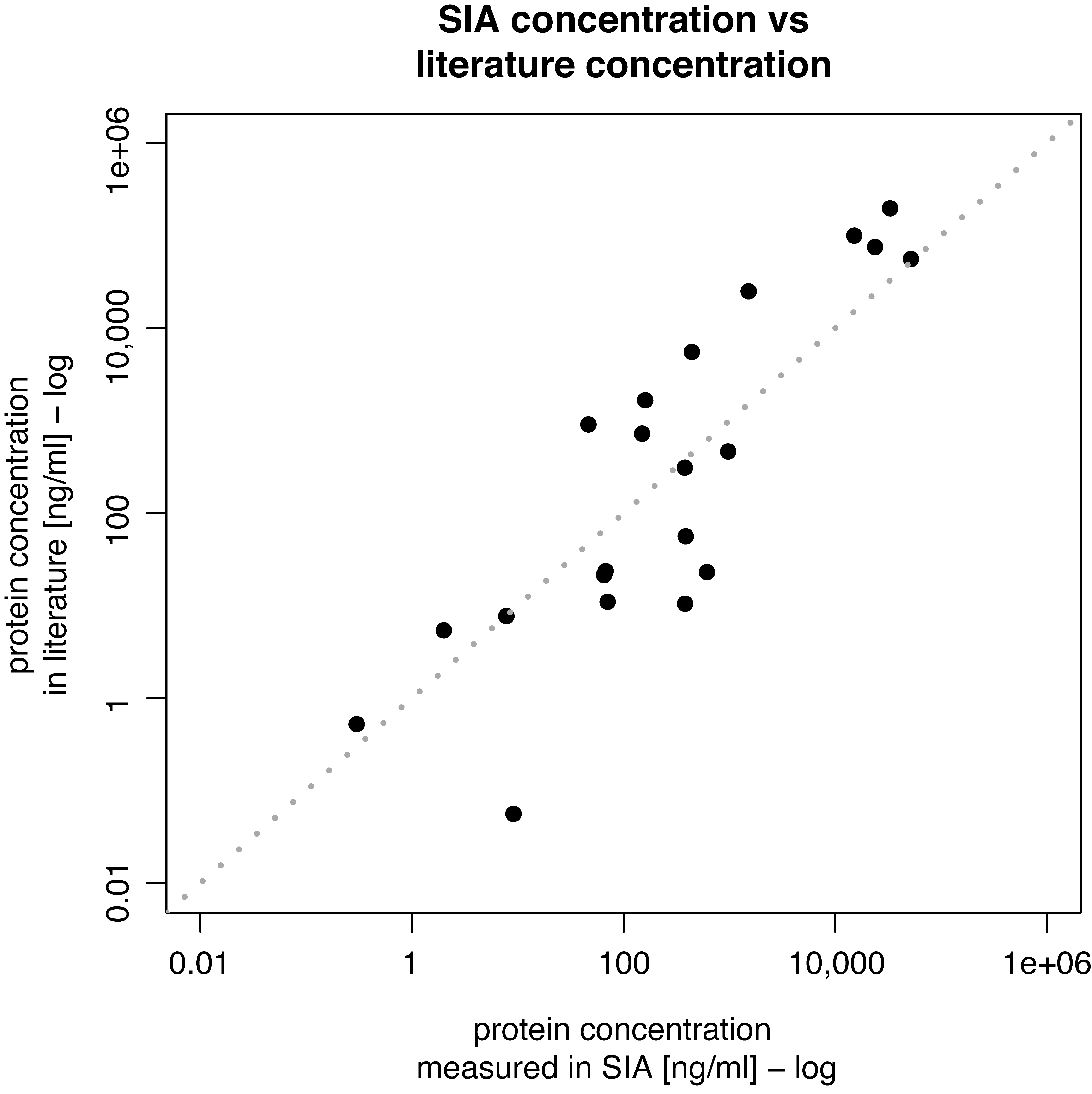
