## Supplementary Figure 3. Functional SIA pairs screening # 1 and # 2 for "Systematic Development of Sandwich Immunoassays for the Plasma Secretome"

| PLASMA \ PROTEINS | 1 | 0 | sum |
|---|---|---|---|
| 1 | **170** | **77** | 247 |
| 0 | **336** | **535** | 871 |
| sum | 506 | 612 | 1118 |

SCREENING 1

| PLASMA \ PROTEINS | 1 | 0 | sum |
|---|---|---|---|
| 1 | **247** | **96** | 343 |
| 0 | **331** | **378** | 709 |
| sum | 578 | 474 | 1052 |

SCREENING 2
